# Overcoming Cisplatin Resistance in 3D Oral Squamous Cell Carcinoma Models via Nanoparticle-Mediated Pt(IV) Drug Delivery

**DOI:** 10.64898/2026.02.22.707247

**Authors:** Ana Belén Griso-Acevedo, Francisco Navas, Natalia Calvo, Victoria Morales, Beatriz Castelo, Javier González Martín-Moro, María José Moran Soto, Raúl Sanz, José Luis Cebrián-Carretero, Rafael A. García-Muñoz, Ana Sastre-Perona

## Abstract

Advanced oral squamous cell carcinoma (OSCC) patients with cisplatin-refractory tumors face a poor prognosis and limited therapeutic options. Cisplatin-based systemic chemotherapy has long been the gold standard despite producing unmanageable adverse side effects and toxicity. Compared to Pt(II)-based analogs, octahedral Pt(IV)-based compounds have demonstrated remarkable potential as antitumor prodrugs. Pt(VI) derivates are chemically inert remaining intact until internalized within cells, demonstrating higher tolerability and selectivity towards cancer cells. In this study we compare antitumoral response implementing primary patient-derived 2D and 3D OSCC *in vitro* models treated with both cisplatin and a novel Pt(IV) compound. We also test the delivery and efficacy of these anticancer drugs *via* novel encapsulation of this Pt(IV) prodrug in the framework of mesoporous silica nanoparticles (Pt(IV)-cov@MSN). Our results show a significant improvement of delivery and cytotoxicity of Pt(IV)-cov@MSN in both cisplatin-responsive and cisplatin-resistant primary patient-derived *in vitro* models. We also show how Pt(IV)-cov@MSN elicits p53-dependent apoptotic cell death superior to that obtained with cisplatin treatment in OSCCs. These findings highlight 3D-primary models as key tools for drug and nanocarrier testing, as well as potential targeted and selective delivery strategies for novel chemotherapeutic agents.

## Introduction

Oral squamous cell carcinoma (OSCC) represents the most common malignancy among head and neck cancers (HNSCCs) and the 12th most diagnosed cancer type[1], whose incidence is still rising. These tumors arise from the epithelium of the oral mucosa and known risk factors include tobacco and alcohol consumption, infection with oncogenic human papilloma virus strains, among others[2]. Locoregionally advanced disease[3][4] and tumors with distant metastasis are associated with poor outcome and limited therapeutic options[3,5]. Treatment of locoregional disease involves surgical excision[6], with postoperative chemo/radiotherapy in patients at risk of recurrence[3,6]. Nevertheless, adjuvant systemic treatment remains the gold standard for advanced disease. Among the chemotherapeutic agents approved for OSCC, cisplatin remains the most widely used for this purpose, showing high antitumor activity and acting as a radiosensitizer in combined chemo-radiotherapy regimens[7]. The cytotoxic effect caused by cisplatin relies on its cellular uptake mainly by copper transporter CTR1[8] among others[9,10]. Cisplatin is then able to bind to nucleophilic centers of the DNA by forming preferentially guanine-base intra- and interstrand adducts. These result in DNA damage, p53 activation and ultimately apoptosis[11,12]. However, platinum-based drug resistance occurs in a large proportion of advanced HNSCCs showing response rates below 40% in patients treated with cisplatin regimens[13,14], representing one of the most common causes of therapeutic failure and recurrence in OSCCs [12]. Systemically administered cisplatin, in addition to other analogs such as carboplatin and oxaliplatin, is associated with important negative side effects, which can also impact the viability of the treatment plan[15].

To overcome these drawbacks, Pt(IV) compounds provide an interesting alternative. These species derive from Pt(II) complexes but differ in various aspects[16,17]. Pt(II) drugs exhibit square-planar geometry (**Fig. 1A**), allowing for four total coordination sites for ligands, arranged around the Pt center[18]. This metallodrug does not require additional reduction and becomes active once administered and therefore, also degraded, reducing bioavailability[19]. Pt(IV) complexes present octahedral geometry, allowing for two additional coordination sites that can be further functionalized, representing a more stable and therefore, inert compounds that do not undergo premature degradation or binding to bloodstream plasma proteins[20], being also resistant to gastric hydrolysis which enables their oral administration[17,21]. Pt(IV) complexes are classified as prodrugs that are integrated into cells through a combination of passive diffusion and active transport depending on ligand composition, with the subsequent additional reduction by different molecules such as glutathione or ascorbic acid within the cell to release the bioactive Pt(II)[16,22,23].

**Fig 1.**
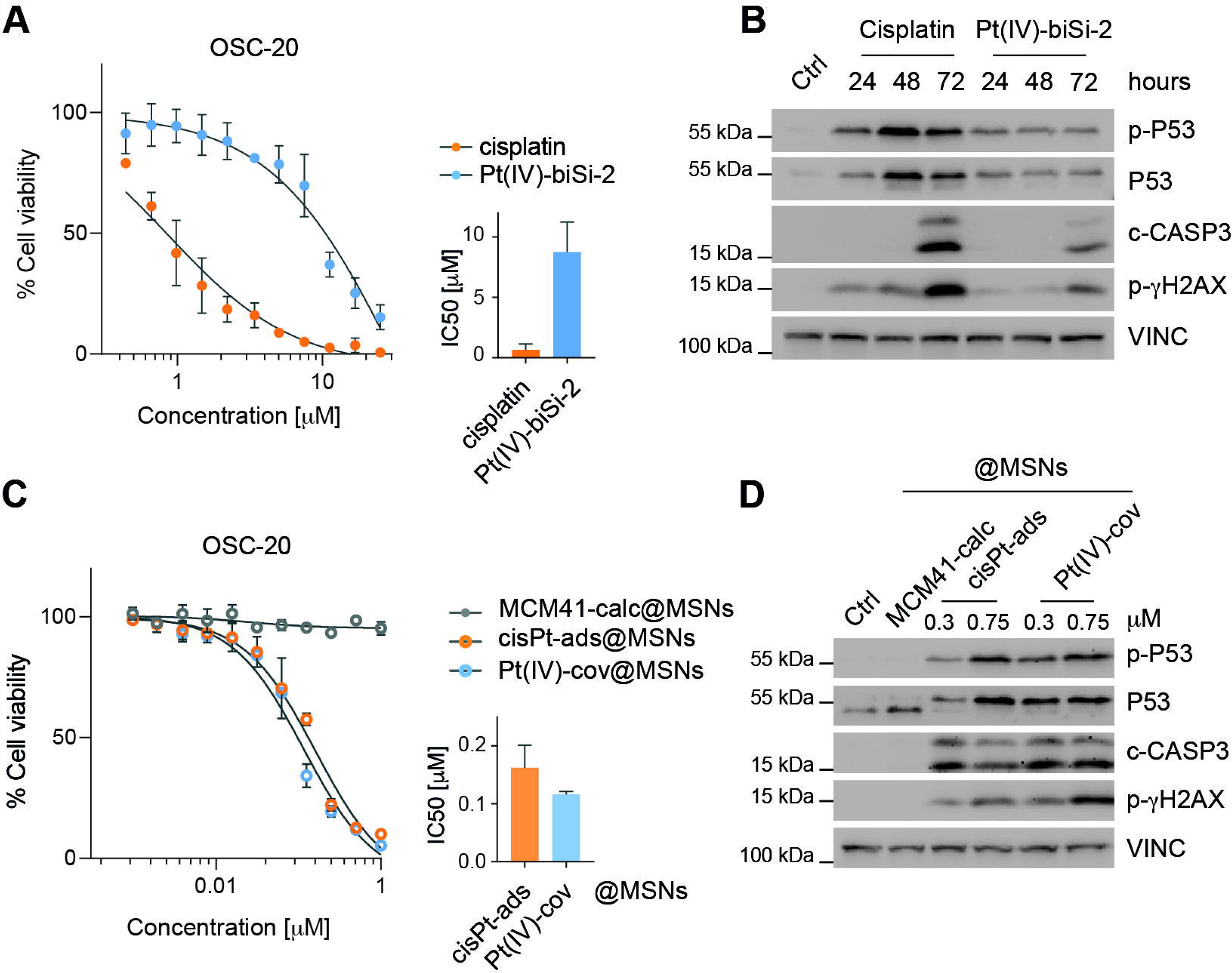
MSN loading of Pt(IV)-biSi-2 improves cytotoxicity in OSCC cell lines. **a)** Dose-response curve for metastatic OSCC cell line OSC-20 with free drug Cisplatin (orange) and Pt(IV)-biSi-2 (blue) concentrations; viability was measured by MTS assay after 7 days. **b)** Apoptotic and DNA damage response markers were assessed by Western Blot analysis of free drug Cisplatin and Pt(IV)-biSi-2 treatment of OSC-20 cells at IC75 concentrations for 24, 48 and 72 h. **c)** Dose-response curve for metastatic OSCC cell line OSC-20 with cisPt-ads@MSNs (orange) and Pt(IV)-cov@MSNs (blue) equivalent concentrations vs. empty control MSNs (MCM41-calc@MSNs, grey); viability was measured by MTS assay after 7 days of treatment. **d)** Western Blot analysis of apoptotic/DNA damage markers after 4 days of IC75 (0.3 µM) and IC90 (0.75 µM) concentration treatments with drug-loaded MSNs after 4 days. Control MSNs (MCM41-calc@MSNs) were added at the highest (IC90) concentration.

Furthermore, emerging strategies like nanocarrier encapsulation of platinum-based compounds aim to ameliorate adverse side effects associated with non-specific cytotoxicity, and improve drug distribution, bioavailability and rapid blood clearance[24–26]. Among these delivery systems, mesoporous silica nanoparticles (MSNs) have gained relevance as nanocarriers in the biomedical field due to their excellent biocompatibility, highly modifiable structure, composition, cargo and functionalization[27–31], demonstrating promising results with platinum-based compounds[24,32]. MSNs have been tested in conventional 2D established cell line monolayers from a wide array of cancer subtypes[33], although these models often are not representative of an *in vivo* tumor in terms of spatial arrangement, cell fate and differentiation, and naturally occurring nutrient, oxygen and treatment gradients present in a patient, thus tend to overestimate drug response[33,34]. To tackle this, 3D cell culture models from patient samples like tumor spheroids or organoids -termed tumoroids- are replacing conventional cell monolayers for drug testing/screening. These models represent the inter- and intra-tumoral heterogeneity that is associated with cisplatin resistance, minimal residual disease and recurrence.

In this study we employ primary, patient-derived 3D OSCC spheroids and organoids to study treatment response, in a comparative and mechanistic analysis of antitumoral cytotoxicity between the classical Pt(II) compound cisplatin and the new Pt(IV) prodrug, termed Pt(IV)-biSi-2 (**Fig. 1A**) which has shown enhanced selectivity and cytotoxicity towards colorectal cancer cells both *in vitro* and *in vivo*[35]. This work evaluates the potential improvement in cytotoxic activity through alternative drug encapsulation of this Pt(IV) prodrug in the three-dimensional structure of MSNs, which release and disperse gradually and protect the metal complexes from premature binding or degradation. To our knowledge, this is the first study using primary 3D OSCC culture models such as spheroids and organoids as drug testing platforms for novel Pt(IV) compounds against conventional cisplatin, both as free drug and nanoparticle formulations.

## Methods

### Chemicals

Hexadecyltrimethylammonium bromide (CTAB ≥ 98%), tetraethyl orthosilicate (TEOS, 98%), (3-aminopropyl)trimethoxysilane (APTMS, 98%), dry toluene (99%), dimethyl sulfoxide (DMSO), phosphate buffered saline (PBS, pH = 7.4), glacial acetic acid (≥99.7%) and sodium acetate anhydrous (>99%) were obtained from Sigma-Aldrich. Hydrochloric acid (35% w/w), sodium hydroxide and absolute ethanol were obtained from Scharlab. Cisplatin (99% Pt) was purchased from Strem Chemicals Inc.

### Clinical Specimens

Fresh human OSCC specimens were obtained from patients at the Hospital Universitario La Paz (Madrid, Spain) under informed consent and approval of the Biobank and Ethical Committees of the hospital according to Spanish ethical regulations. The study followed the guidelines of the Declaration of Helsinki, and patient identity and pathological specimens remained anonymous in the context of the study.

### Cell culture

Patient-derived OSCC primary cultures were obtained from surgical discards provided by the La Paz University Hospital Oral and Maxillofacial surgical team were established following a primary tissue-processing protocol modified from Schober et al.[36]. Tumoral tissue was mechanically shredded and digested with 2.5 mg/mL Collagenase I (Sigma-Aldrich) in Advanced+++ medium (adDMEM/F12 medium (Gibco) with 1X GlutaMAX^TM^ (Gibco), 1X Penicillin/Streptomycin (Corning) and 10 mM HEPES (Gibco)) with Rock-inhibitor 10 µM (Bio-techne) for 45 minutes at 37°C shaking. DNAse I (Roche) was added for 10 additional minutes, before cell suspension was retrieved with Ca2+/Mg2+-free PBS (Biowest) and filtered through 70 and 40 µm cell strainers (Corning). Tissue was retained on filters and further digested with 0.25% Trypsin (Gibco) for 10 minutes at 37°C in an orbital shaker, before it was strained and combined with prior isolated. Digested mix was centrifuged at 450 g 10 min at 4°C. Red blood cells were removed by resuspending pellet in 1X RBC buffer (Miltenyi Biotec) and centrifuged 450 g 4°C for 5 min before seeding cells on a Mitomycin C (#10107409001, Roche) treated 3T3-J2 murine embryonic fibroblast monolayer. Digested epithelial cells were seeded with E-medium (DMEM/F12 3:1 (Corning), 10% FBS, 1X Penicillin/Streptomycin, 1X GlutaMAX™, 7.5 mM HEPES buffer, 3.75 ug/mL Apotransferrin, 15 pM T3 hormone, 3.75 µg/mL Insulin, 30 ng/mL Hydrocortisone and 7.5 pM Cholera toxin (Sigma-Aldrich)), supplemented with 10 µM Rock inhibitor and 50 ng/mL hEGF (Peprotech), as well as 50 µg/mL Primocin^TM^ (Invivogen) and 1 µg/mL Caspofungin (Sigma-Aldrich). For further experiments, murine feeder cells were removed and remaining epithelial colonies were washed with 1X PBS and treated with trypsin 0.25% to obtain enriched tumoral single cell suspensions.

### Spheroid generation and treatment

To generate organoids, 5,000 cells were seeded in FGF-10 supplemented (10 ng/mL) OncoPro™ (Gibco) medium. 3% BME (Bio-Techne) was added to the cell-medium mix and seeded in 96-well ULA (Corning) plates, centrifuged at 4°C 1500 rpm for 10 min and left 72h hours to assemble and grow before treatment. Cells were treated with soluble cisplatin and Pt(IV)-biSi-2 to a final concentration of 10 µM. As for the platinum-based nanoparticles, these were sonicated and a volume equivalent to 10 µM of cisplatin or Pt(IV)-biSi-2 cargo was calculated and added to the spheroid medium. Treatment was sustained and monitored for 7 days before further downstream applications.

### BME-embedded organoid generation and treatment

To generate BME-embedded organoids, 50,000 cells per 50 µL drop were seeded in 70% BME/ 30% Advanced +++ mix in p24 ULA well plates. FGF-10-supplemented OncoPro™ medium was used to overlay the BME droplets 72h before treatment. Both soluble drugs and platinum-based MSNs were added to a final concentration of 10 µM in fresh FGF-10-supplemented OncoPro™ medium for 7 days before further downstream applications.

### Immunohistochemistry

Organoids were fixed by removing the overlaying medium and adding 4% PFA (#28908, Thermo Scientific) for 30 min at RT, retrieving the released organoids carefully with a cut 1000-pipette tip in a 1.5 mL tube. Organoids were centrifuged at 300 g for 5 min and washed once with 1X PBS before serial dehydration. The organoid pellet was resuspended in 70°, 80°, 96° and 100° ethanol for 15 min each, centrifuging at 300 g 5 min in between each ethanol incubation, and then resuspended in xylene for 30 min. Once xylene supernatant is removed and pellet is left to air-dry as much as possible, hot paraffin was added up to halfway the 1.5 mL tube and kept at 60° C in a thermoblock or water bath for 1 h before letting the paraffin-embedded pellet cool and solidify within the tube. The end tip of the tube -containing the pellet- was sliced and contents were mounted in paraffin blocks for further sectioning. Hematoxylin-Eosin staining was performed on treated BME-organoid 5 µm sections after 30 min deparaffinization at 60°C, following a standard protocol for this procedure. Immunohistochemistry DAB staining was performed for 5 µm organoid sections which were dewaxed for 1 h at 60°C and then rehydrated taking the following steps: Xylene for 6 min, 100° EtOH for 6 min, 95° EtOH 6 min, 70° EtOH for 3 min, 50° EtOH for 3 min and leaving the slides in distilled water. Antigen retrieval was performed by incubating the rehydrated slides in a DAKO PT LINK pre-treatment module previously filled with pre-heated Tris-HCL buffer (pH=9) at 90°C for 20 min. Slides are again left in dH20 until adding BLOXALL® (#SP-6000-100, Vector Laboratories) for 10 min to block endogenous peroxidase activity and then washing with PBS for another 5 min. Excess liquid was removed to draw on individual chambers with a hydrophobic or PAP-pen (#101098-065, Vector Laboratories) before blocking. Tissue was then incubated for 15 min at RT with Avidin/Biotin-supplemented Goat normal blocking serum (#SP-2001, #AK-5001, Vector Laboratories) and washed again with PBS. Slides were incubated incubate at 4°C O/N with primary antibody diluted in blocking buffer. Slides were then washed 3 times with PBS and incubated with biotinylated secondary antibody diluted 1:200 in blocking serum (#AK-5001, Vector Laboratories) for 1 h at RT before washing twice with PBS. Tissue was then stained with DAB (#SK-4105) for 10 min before rinsing with water, counterstaining with hematoxylin, rinsing, dehydrating again (following the previous rehydration steps in the opposite order) and mounting with DPX mounting medium). Primary antibodies: anti-Cleaved Caspase-3 (#9664, Cell Signaling, 1:1000), anti-P53 (#9282, Cell Signaling, 1:1000). IHQ pictures were acquired at 20X magnification using Image-Pro software using a brightfield microscope.

### Organoid Immunofluorescence

BME-embedded organoids were released and retrieved from the BME by digestion with 2 mg/mL Dispase® II (#D4693, Sigma-Aldrich) 1:50 in Advanced +++ medium supplemented with 10µM Rock inhibitor 30 min at 37° C. One dissolved, organoids were retrieved using a cut 1000-pipette tip and centrifuged in 1.5 mL tubes at 300 g for 5 min. Organoid pellet was washed twice with ice-cold 1X PBS to remove and dissolve excess BME. Organoids were then fixed overnight at RT with 4% formaldehyde and pelleted again. Fixating agent was discarded and organoid pellet was resuspended with 150 −200 µL of preheated, liquid 4% noble agar (#A5431, Sigma-Aldrich) and mix was left to solidify at RT in a 7 x 7 x 5 mm stainless steel embedding base mold. Organoid-embedded agar squares were sequentially dehydrated in 70°, 80°, 96°, 100° EtOH and xylene for 1 h each before paraffin embedding. 5 µm sections were dewaxed at 60 °C for an hour and rehydrated by sequentially incubating in: xylene (10 min), 100° EtOH (10 min), 96°, 80° and 50° EtOH (5 min each) before leaving slides in distilled water. Antigen retrieval was performed by incubating the rehydrated slides in a DAKO PT LINK with Tris-EDTA buffer (pH=9) at 90°C for 20 min. Tissue sections were blocked for 1h at RT with blocking buffer (1% BSA, 5% Goat Serum, 0,3% Triton X-100 in 1x PBS). After blocking buffer was removed, and tissue incubated with primary antibody diluted in blocking buffer at 4° C overnight. After three washes with 1X PBS tissues were incubated with secondary antibody and DAPI diluted in blocking buffer for 1 h at RT. Tissues were again washed thrice with 1x PBS and mounted with ProLong™ Diamond (#P36970, Invitrogen) mountant. Images were taken at 20X magnification using a Leica STELLARIS 8 spectral confocal microscope. Primary antibodies: anti-KI67 (#RM9106-50, Epredia, 1:400), anti-P63 (#13109, Cell Signaling, 1:1000). Secondary antibodies: Donkey anti-Rabbit Rhodamine Red X (#JAC-711-295-152, Jackson Immunoresearch, 1:1000), DAPI (#D1306, Invitrogen, 1:1000).

### Annexin V/7AAD staining

Medium containing spheroids was retrieved together from wells using a cut 200-pipette tip and centrifuging at 300 g for 5 min, before setting on ice. For BME-embedded organoid retrieval, BME was digested with 2 mg/mL Dispase® II (#D4693, Sigma-Aldrich) 1:50 in Advanced +++ medium supplemented with 10µM Rock inhibitor 30 min at 37° C. One dissolved, organoids were retrieved using a cut 1000-pipette tip and centrifuged at 300 g for 5 min before setting on ice. Pellet (both from spheroids and organoids) was washed twice with ice-cold serum-free medium and incubated in ice for 20 min, followed by a centrifugation at 300 g 5 min at 4°C. Pellet was resuspended TrypLE™ Express (#12604013, Gibco™) and mechanically disrupted using a 1000-10 µL compound tip before incubating for 10 min at 37°C, pipetting with compound tip periodically. FBS was added at 20% final volume to stop trypsin digestion and cells were visualized through a hemocytometer to check for single cell suspension. Cells were centrifuged again and resuspended in 1X Binding Buffer for further Annexin V and/or 7AAD staining (Annexin V-FITC Kit #130-092-052, 7-AAD Staining solution #130-111-568, Miltenyi Biotec). After centrifugation for 10 min at 200 rcf, cells were resuspended in staining solution consisting of diluted Annexin V and/or 7-AAD antibodies in 1X Binding Buffer and incubating for 15 min at RT, protected from light. 150 µL of 1x Binding Buffer was added to further dilute the staining solution before analyzing samples employing a BD FACSCelesta™ multicolor flow cytometer.

### Cell Cycle staining

Single cell suspension from either spheroids or organoids was obtained following the steps described above (See Annexin V/7AAD staining), centrifuged and resuspended in FACS buffer (1X Ca2+, Mg2+-free PBS, 3% FBS, 0.06% DNAse I) and stained with Hoechst (#H3570, Invitrogen^TM^) 1:1000 dilution for 45 minutes at 37° C, resuspending periodically. Samples were further diluted with FACS buffer and analyzed employing a BD FACSCelesta™ multicolor flow cytometer.

### 3D culture viability analysis /ATP measurement

3D cultures (organoids and spheroids) were collected using a cut 200 tip on a micropipette and incubated with equal volume of RT Advanced +++ medium and CellTiter-Glo® 3D Reagent (#G9681, Promega) for a total of 30 minutes, protected from light, at RT. The first 5 and last 5 minutes were used to induce cell lysis by mixing in an orbital shaker (25°C, >1000 rpm). The mix was then pipetted in opaque p96-well plates and luminescence values were acquired using Promega GloMAX® Discover plate reader using an integration time of 0.3 seconds per well.

### Dose-response curves

For the dose-response (DR) curves, 5,000 cells per well were seeded in 96-well plates (#167542, Thermo Scientific) in their appropriated media. 24 h later, cells were treated by performing 12 sequential, 2/3 serial dilutions, ranging from 25 to 0 µM. Cells were incubated with the drugs for 7 days and MTS assay (#G5430, Promega) was performed to analyze cell viability following the manufactures instructions and data was acquired in an Epoch (BioTek Instruments) Microplate Reader at 490 nm. For MSN-loaded drug dose response curves, resistant cell lines were treated with 0 to 25 µM concentration using 2/3 serial dilutions, whereas sensitive cell cultures were treated with 0-1 µM concentration with 1/2 serial dilutions. Empty control MSNs were added in equivalent concentrations as toxicity control.

### Western Blotting

Protein extraction was performed on ice, washing with 1X PBS, and adding RIPA lysis buffer (150 mM NaCl, 1% Triton X-100, 0.5% Sodium deoxycholate, 0.1% SDS, 50 mM Tris pH=8, diluted in distilled water) supplemented with protease and phosphatase inhibitors (#05892970001, #PHOSS-RO, Roche). Cell lysates were incubated 10 minutes on ice and then centrifuged at 12.000 g for 10 min at 4°C to collect protein supernatant and quantify concentration by performing Bradford protein quantification assay (#5000006, Bio-Rad). SDS-PAGE Western Blotting was performed using 20-30 ug of total protein per condition in 12% polyacrylamide gels. After gel transfer to nitrocellulose membranes, these were stained with 0.1% Ponceau in 5% acetic acid and further blocked in 5% skimmed milk powder in TBST (1X TBS; 0.05% Tween) for 45 minutes at RT. Primary antibodies were diluted in either 5% Skimmed milk powder or BSA in TBST and incubated O/N at 4°C on a shaker. Primary antibody was retrieved, and membranes were washed 3x with TBST before incubating in secondary HRP-conjugated antibodies for 1h at RT. Membranes were then again washed and incubated for 1 min in chemiluminescent substrate ECL (34580, Thermo Scientific) before imaging with an UVITEC ALLIANCE imager. Primary antibodies: anti-Cleaved Caspase-3 (#9664, Cell Signaling, 1:1000), anti-p-γH2AX (#2577, Cell Signaling, 1:1000), anti-p-P53 (#9284, Cell Signaling, 1:1000), anti-P53 (#9282, Cell Signaling, 1:1000), anti-Vinculin (#sc-73614, Santa Cruz, 1:1000). Secondary antibodies: anti-Rabbit HRP (#A27036, Invitrogen, 1:10.000), anti-Mouse HRP (#AP124P, Sigma-Aldrich,1:1000-2000).

### Image Processing

Brightfield 4X images were acquired using a Leica DMI6000B microscope and preprocessed with FIJI ImageJ software. Organoid images were segmented using a size pixel-based classification algorithm within Ilastik software[37] and further processed for size measurement using FIJI ImageJ software.

### Data and Statistical Analysis

Comparative data is presented as mean (±SEM) of at least 3 biological experiments. Statistical analysis was performed using GraphPad Prism v.9.1.0. Two-tailed Student’s *t* tests were used for comparisons between two conditions; for more than two groups ANOVA and Tukey’s test were used and significant differences are indicated with asterisks. Differences are considered significant at *p*□<□0.05 (*), *p*□<□0.01 (**), *p*□<□0.001 (***), *p*<0.0001 (****).

### Illustrations

Schematic illustrations were created using Adobe Illustrator.

## Results

### Encapsulation of Pt(IV)-biSi-2 in MSNs improves efficacy in metastatic OSCC cell line OSC20

To compare the cytotoxicity of cisplatin and Pt(IV)-biSi-2, we used the OSC-20 cell line, which was derived from a metastatic tongue squamous cell carcinoma. OSC-20 cells showed sensitivity to cisplatin in MTS assay, measuring a rapid decrease in cell viability upon treatment with increasing dosage of cisplatin showing an IC50 of 0.8 µM. When treated with identical increasing concentrations of Pt(IV)-biSi-2, these cells exhibited decreased sensitivity, showing an 8 µM IC50, almost ten-fold the one obtained with cisplatin (**Fig. 1A**). Since this novel Pt(IV) prodrug had not yet been tested on OSCC cell line models, we analyzed whether observed cell death with this compound occurred through p53 activation. Cells were treated with IC75 concentrations of both drugs at different times and collected for Western Blot analysis (**Fig. 1B**). Interestingly, DNA damage was already present at 24h for both drugs, with increasing intensity of p-γH2AX in consecutive timepoints. This also correlated with P53 accumulation and activation (p-P53) over time, resulting in an increase of caspase-3 cleavage. Although Pt(IV)-biSi-2 treatment-induced activation of DNA response and p53 activation, the intensity seems to be weaker and delayed in time. Ultimately, caspase-3 cleavage was also observed at the 48h mark, indicating apoptotic cell death, although this was decreased in comparison with cisplatin.

We tested the effect of encapsulated cisplatin and Pt(IV)-biSi-2 in MSNs (cisPt-ads@MSNs and Pt(IV)-cov@MSNs, respectively) which would gradually release the drugs for 7 days. We recapitulated the dose-response experiments, while adding empty vehicle nanoparticles (MCM41-calc@MSNs) as a control. To our surprise, Pt(IV)-cov@MSNs displayed an improved cytotoxic effect on OSC-20 cells than the soluble version, reducing IC50 values from 8.75 µM to 0.1 µM (**Fig. 1A,C**). cisPt-ads@MSNs slightly lowered its IC50 value when administrated in this format (from 0.65 µM to 0.2 µM) and unexpectedly double the IC50 of Pt(IV)-cov@MSNs. When cells were treated at IC75 and IC90 concentrations with MSN-encapsulated drugs (**Fig. 1D**), we observed higher activation and accumulation of p-P53 and total P53 with Pt(IV)-cov@MSNs at lower concentrations. At higher IC90 concentrations, Pt(IV)-cov@MSNs also induced more DNA-damage which concluded in caspase-3 cleavage. These results demonstrate that as cisplatin, Pt(IV)-biSi-2 induces p53 and caspase-3-dependent cell death in OSCC cell line OSC-20, and this is greatly improved by MSN encapsulation, where Pt(IV)-biSi-2 elicits greater cytotoxic effect than cisplatin.

### MSN-encapsulated Pt(IV)-biSi-2 outperforms cisplatin in cisplatin-responsive primary patient-derived OSCC cell lines

These results encouraged further testing in more biologically relevant and translational models such as primary, patient-derived 3D OSCC models. We establish primary cultures from stage III-IV OSCC tumors[36]. To test their sensitivity to cisplatin, these cultures were grown in BME into patient-derived organoids [38] before treatment with cisplatin. OSCC43 organoids showed stalled organoid growth after treatment in comparison to OSCC37 and OSCC55 organoids, which continued growing and showed minimal growth stunting after treatment (**Fig. 2A,B**). Therefore, we classified the organoids into cisplatin-sensitive (OSCC43) or resistant (OSCC37 and 55) and used them to further test cisplatin and Pt(IV)-biSi-2 formulations. Cisplatin-responsive OSCC43 cells were appropriate candidates for 2D and 3D testing, since they were the most proliferative, as seen through proliferative marker KI67 staining of control organoids (**Fig. 2C**). Cisplatin-resistant samples form stratified and differentiated organoids which are less proliferative, form keratinized cores markedly visible through eosin staining (**Fig. 2B**) while maintaining an outer basal epithelial stem cell population positive for marker P63 (**Fig. 2C**).

**Fig 2.**
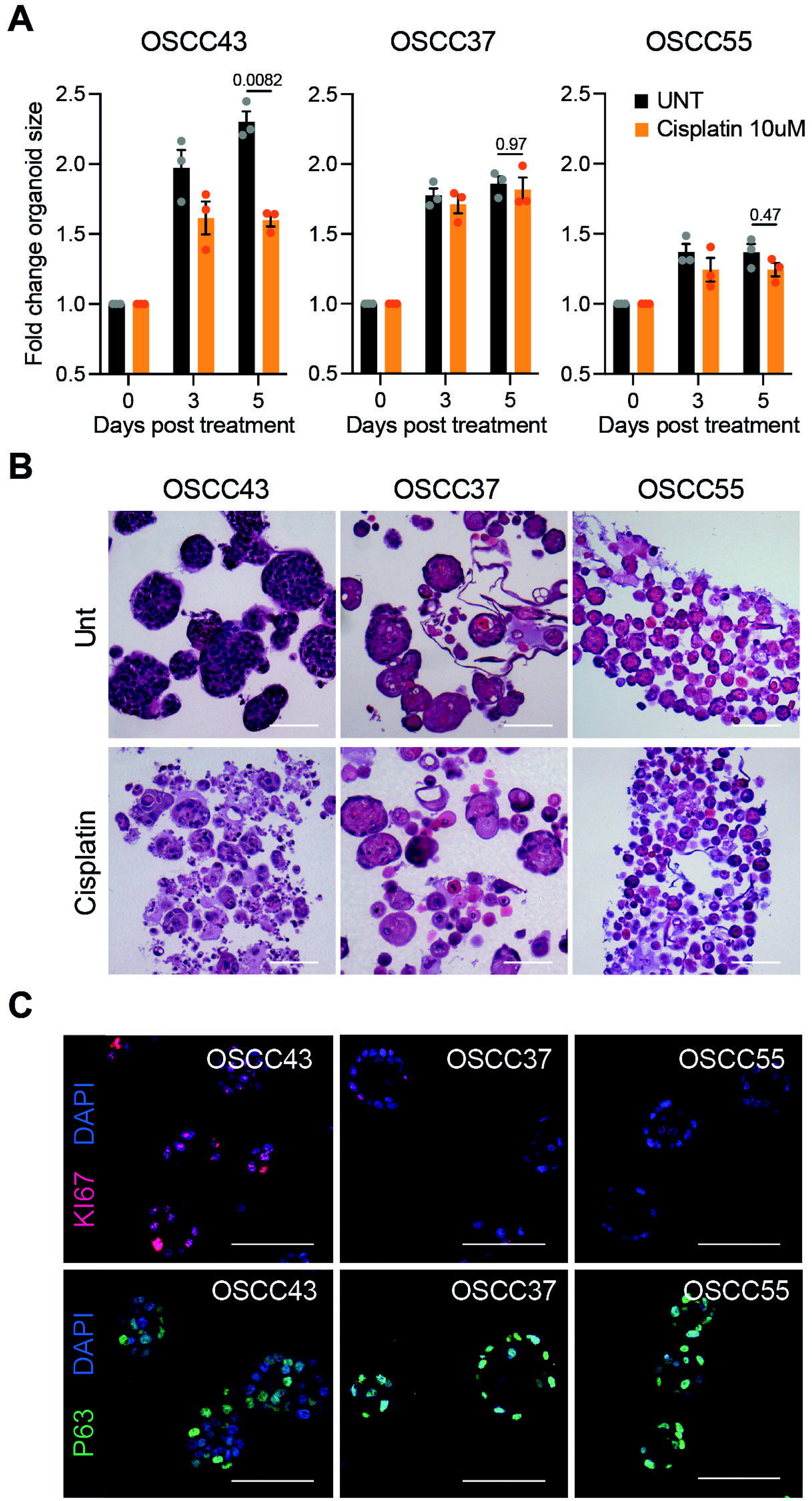
Characterization of OSCC organoids in response to cisplatin. **a)** Fold change of mean organoid area between endpoint and start of treatment of OSCC primary tumor-derived samples OSCC43, OSCC37 and OSCC55 grown in BME. Organoids were grown for 3 days prior to adding 10 µM of soluble cisplatin for 5 days (displayed in orange) or left untreated (UNT; dark grey). **b)** Hematoxylin-Eosin staining of paraffin-embedded organoids fixed at endpoint depict organoid size at 5 days between treated and untreated conditions. **c)** Immunofluorescence of untreated organoids for nuclear staining of KI67 (upper row, red) and P63 (bottom row, green). Nuclei were stained with DAPI (blue). Scale bar = 100 µm

Cisplatin dose-response MTS analyses (**Fig. 3A**) confirmed a drastic decrease in cell viability cisplatin (0.6µM IC50), whereas cell viability reduction with increasing doses of Pt(IV)-biSi-2 was detected at tenfold higher concentrations (6µM IC50). Western Blot analysis of cells treated with IC75 concentrations of the drugs showed that the Pt(IV)-biSi-2 induced a lagged activation of P53 and caspase-3 cleavage (**Fig. 3B**), confirming what was observed in the OSC-20 cell line.

**Fig 3.**
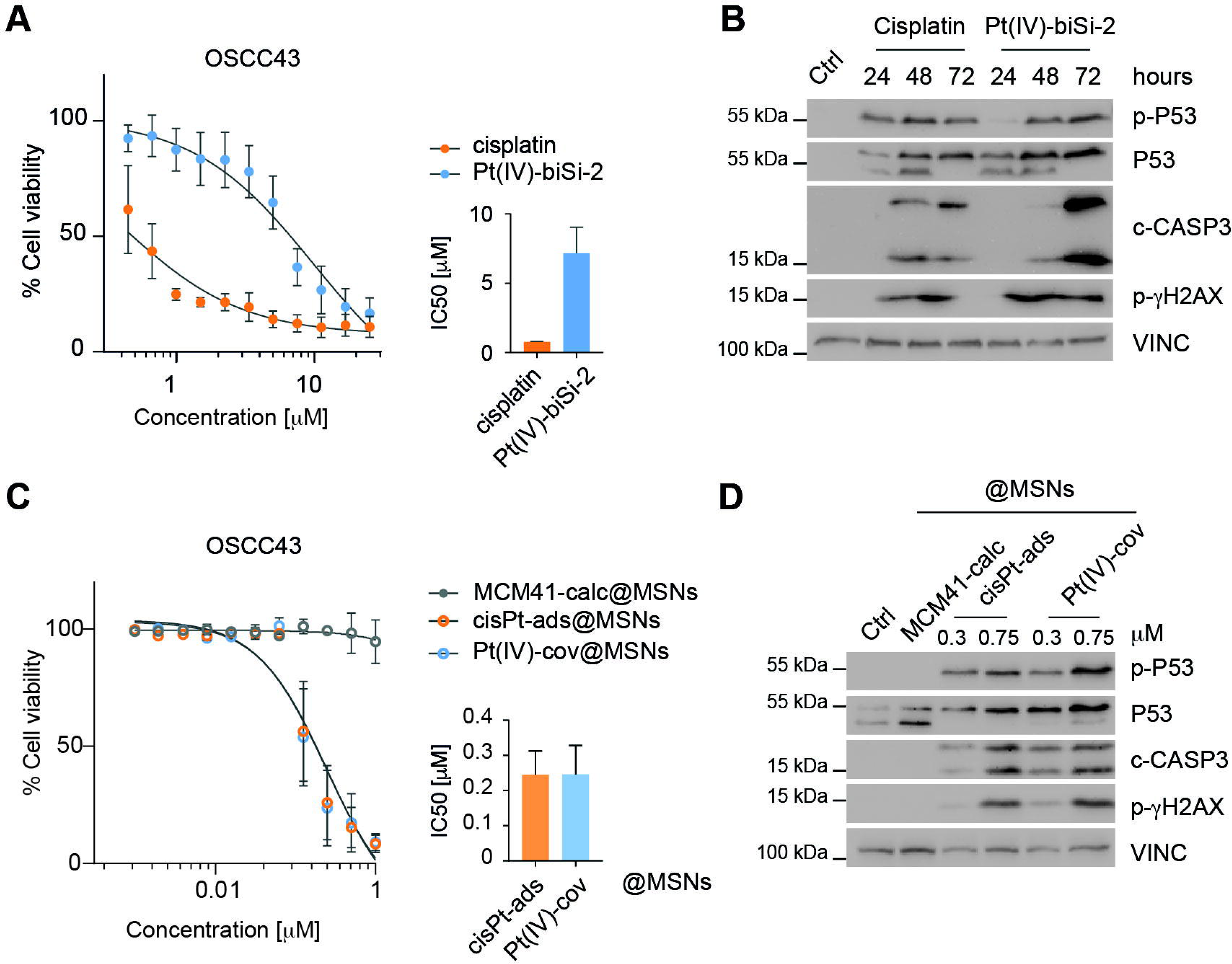
MSN loading of Pt(IV)-biSi-2 improves cytotoxicity of OSCC-sensitive primary cultures. **a)** Dose-response curve for primary cisplatin-sensitive OSCC43 cell line with free drug Cisplatin (orange) and Pt(IV)-biSi-2 (blue) concentrations; viability was measured by MTS assay after 7 days. **b)** Apoptotic and DNA damage response markers were assessed by Western Blot analysis of free drug Cisplatin and Pt(IV)-biSi-2 treatment of OSCC43 cells at IC75 concentrations for 24, 48 and 72 h. **c)** Dose-response curve for OSCC43 cell line with cisPt-ads@MSNs (orange) and Pt(IV)-cov@MSNs (blue) equivalent concentrations vs. empty control MSNs (MCM41-calc@MSNs, grey); viability measured by MTS assay after 7 days of treatment. **d)** Western Blot analysis of apoptotic/DNA damage markers after 4 days of IC75 (0.3 µM) and IC90 (0.75 µM) concentration treatments with drug-loaded MSNs. Control MSNs (MCM41-calc@MSNs) were added at the highest (IC90) concentration.

Next, we tested the effect of loading the drugs in MSNs (**Fig. 3C**). Again, we observed a drastic reduction in IC50 values for Pt(IV)-cov@MSNs, as well as a moderate reduction for cisPt-ads@MSNs (7 µM and 0.75 µM, respectively to 0.2 µM). When we treated the cells at IC75 a IC90 for protein analysis, Pt(IV)-cov@MSNs elicited higher accumulation of p-P53 and total P53 than equivalent concentrations of cisPt-ads@MSNs, as well as an increase in caspase-3 cleavage at IC75. DNA damage shown through p-γH2AX induction was clearly visible at IC90 concentrations. Taken altogether, these results corroborate the earlier hypothesis that Pt(IV)-biSi-2 elicits a greater P53, caspase-3 dependent apoptotic process than cisplatin in cisplatin-responsive OSCC cell lines, which is facilitated by MSN encapsulation.

### Primary patient-derived 3D *in vitro* models show effective integration of MSNs and increased cytotoxicity of Pt(IV)-cov@MSNs

Cisplatin-responsive primary line OSCC43 was used to generate two 3D models for *in vitro* testing of soluble and MSN-encapsulated cisplatin and Pt(IV)-biSi-2. Cells were grown both into single spheroids per well with a low BME concentration (3%) (**Fig. 4A**), and organoids in highly concentrated (70%) BME matrices (**Fig. 4B**). Spheroids and organoids were subsequently treated with 10 µM of each compound before further downstream processing. Measurement of the relative final spheroid and organoid size showed a significant decrease with cisplatin treatment, most pronounced in spheroid models (**Fig. 4A-D**), while Pt(IV)-biSi-2 restricted spheroid growth, although this effect was also less evident in BME-embedded organoids. Since the cytotoxic effect of both free compounds appeared to be attenuated in BME-embedded organoids, we tested if the matrix might interfere with compound absorption, decreasing MSN-encapsulated drug uptake. We therefore conducted an integration assay with FITC-conjugated MSN-encapsulated compounds in both models, which were treated and disaggregated into single cell suspension for flow cytometry analysis (**Fig. 4E**). All three nanoparticle formulations integrated effectively into live cells from spheroids and organoids demonstrating that MSN-encapsulated drugs reached organoids despite being embedded in BME matrices, which was also observable in hematoxylin-eosin-stained organoid sections (**Fig. S1A**).

**Fig 4.**
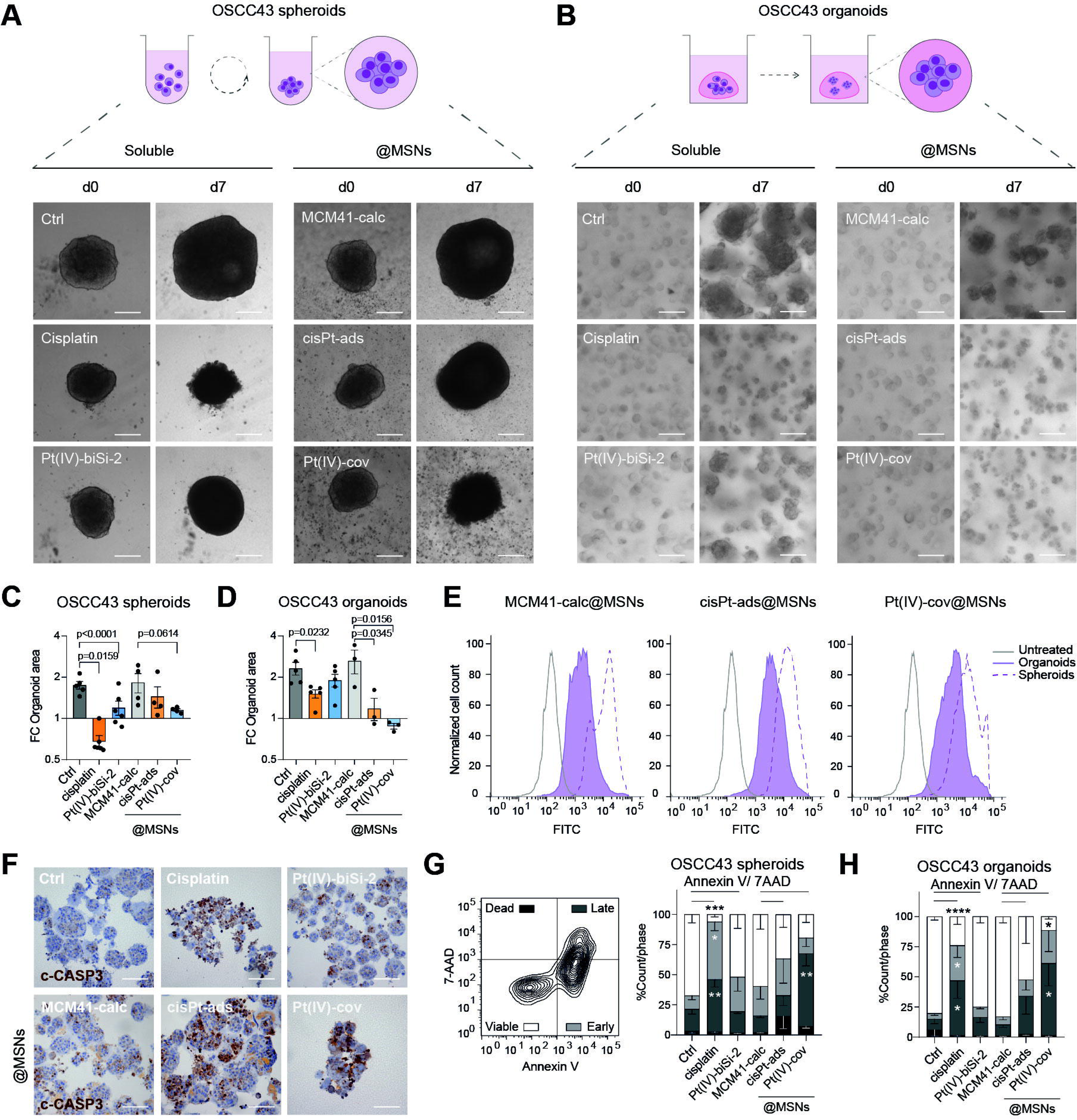
MSN loading of Pt(IV)-biSi-2 improves cytotoxicity of OSCC-sensitive 3D primary models. Cisplatin-responsive primary cell line OSCC43 was grown in two 3D *in vitro* models for 3 days prior to treatment for 7 days with 10 µM of respective drug combinations, either in free drug formulation or drug-loaded MSNs. Brightfield microscopy was used to capture spheroids (**a**) and BME-embedded organoids (**b**) at beginning of treatment (d0) and at treatment endpoint (d7). (**c**, **d**) Graph showing spheroid and organoid growth throughout treatment expressed as Log2 of fold change in area at day 7 vs day 3, for OSCC43 spheroids and BME-embedded organoids. mean ± SEM, n = 3. Statistical significance is shown as p-value scores. **e**) Representative histograms of flow cytometry analysis of FITC-labeled nanoparticles (MCM41-calc@MSNs, cisPt-ads@MSNs and Pt(IV)-cov@MSNs) within live cells from BME-embedded organoids (BME) and spheroids (SPH) vs an unstained (UNS) control. **f**) Representative images of Caspase-3 Immunohistochemistry of treated OSCC43 organoids. Scale bar = 100 µm. **g**) Left panel, representative flow cytometry chart illustrating 7-AAD and Annexin V staining of treated OSCC43 cells. **g, h**) Graphs showing cell percentage of viable, early and late apoptotic and dead cells from OSCC43 spheroids (**g**) and BME-embedded treated organoids (**h**). mean ± SEM, n = 3, statistical significance is displayed as follows: * = p<0.05; ** = p<0.01; *** = p<0.001; **** = p<0.0001.

In 3D cultures, Pt(IV)-cov@MSNs improved biological response over the soluble version and cisPt-ads@MSN both in spheroid and organoid models (**Fig. 4C-D**). Like previous 2D results, cisPt-ads@MSN displayed a lower activity than in free formulation while MCM41-calc@MSNs control did not have effects on 3D model growth. To confirm the effect of MSN-encapsulated drugs in 3D models, we measured apoptosis through immunohistochemistry staining for cleaved caspase-3 in BME-embedded organoids (**Fig. 4F**), which showed a clear accumulation of activated-caspase-3 more notably in drug-loaded MSNs. This was also confirmed in a quantitative way by flow cytometry with 7AAD/Annexin V-stained of cells from treated spheroids and organoids (**Fig. 4G,H**). This revealed that Pt(IV)-cov@MSNs treatment produced the highest percentage of early and late apoptotic stages in both models. Despite the decrease in organoid/spheroid size in Pt(IV)-biSi-2-treated conditions, Annexin V/7AAD data showed no induction of apoptosis. Additional cell cycle analysis demonstrated that Pt(IV)-biSi-2 did not produce a DNA-damage-induced cell cycle arrest (**Fig. S1B**), however, a significant increase in subG1-cells was observed for cisplatin and drug-loaded-MSNs confirming our previous data. Altogether, these data suggest that Pt(IV)-cov@MSNs elicit better cytotoxic effect in 3D cisplatin sensitive primary models of OSCC through caspase-3-dependent apoptosis.

### Primary patient-derived cisplatin-resistant models show increased response to Pt(IV)-cov@MSNs

Next, we tested the potential use of Pt(IV)-biSi-2 compound on OSCC-cisplatin-resistant models to improve their treatment response. Resistant primary cell lines OSCC37 and OSCC55 (**Fig. 2A-C**) were treated with serial dilutions of cisplatin and Pt(IV)-biSi-2, before performing MTS viability assay (**Fig. 5A, C**). These results confirmed the decreased sensitivity of these resistant cells to both compounds, and the advantage of cisplatin in comparison to Pt(IV)-biSi-2 in their soluble versions (**Fig. 5B, D**). Western blot analysis of treated cells showed that apoptotic caspase-3-dependent cell death was induced as early as 24 h in cisplatin treated OSCC37 and OSCC55 lines while strikingly, no apoptotic death or DNA damage response markers were observed with free Pt(IV)-biSi-2 even after 72 h of treatment.

**Fig 5.**
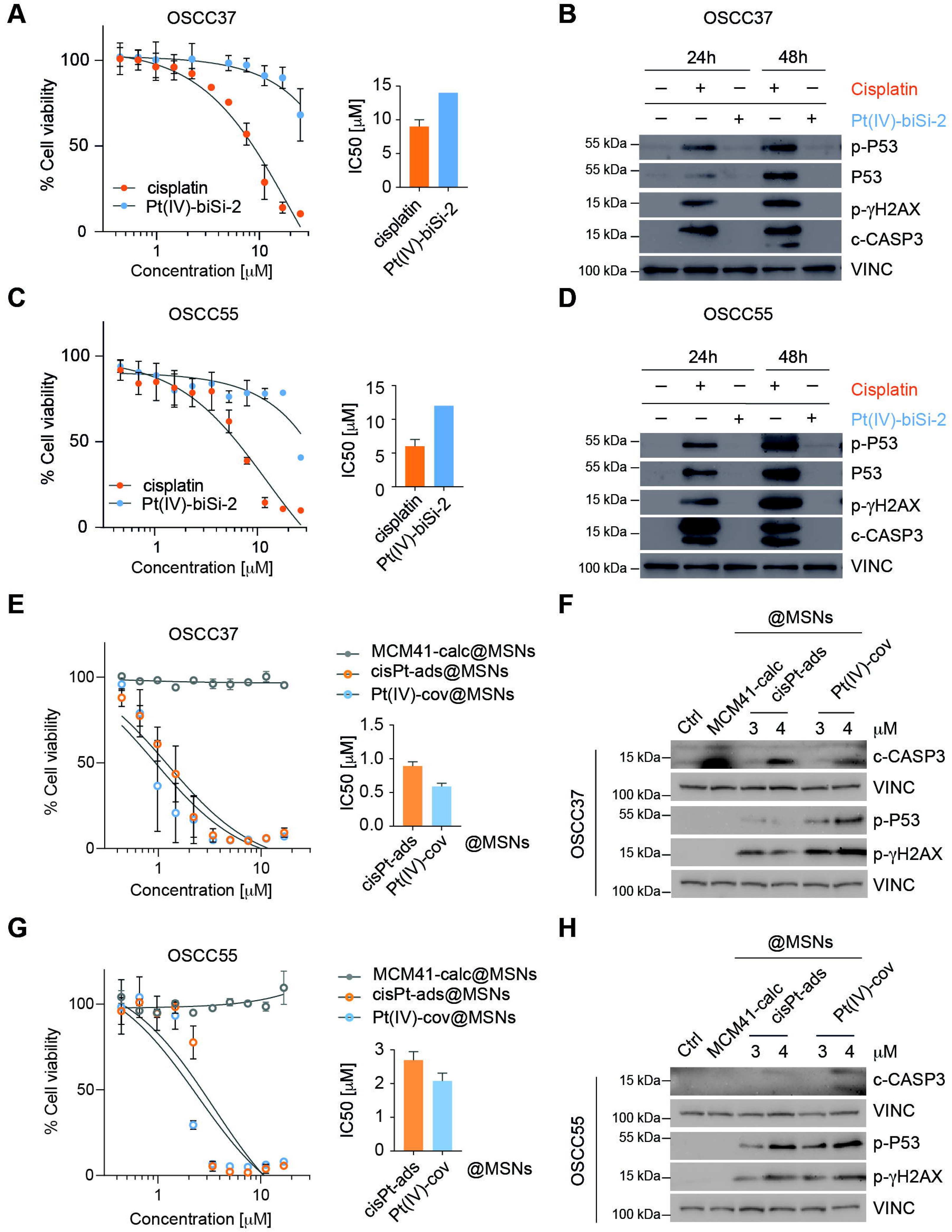
MSN loading of Pt(IV)-biSi-2 sensitize cisplatin-resistant OSCC 2D cultures. Dose-response curve for cisplatin-resistant cultures OSCC37 (**a**) and OSCC55 (**c**) treated with increasing cisplatin (orange) and Pt(IV)-biSi-2 (blue) concentrations (maximum concentration: 25 µM); viability was measured by MTS assay after 7 days. (**b**, **d**) Western Blot analysis for DNA-damage response markers for OSCC37 and OSCC55 cultures treated with maximum 25 µM of cisplatin and Pt(IV)-biSi-2 for 24h and 48h. Dose-response curve for cisplatin-resistant cultures OSCC37 (**e**) and OSCC55 (**g**) treated with MSN-encapsulated drugs cisPt-ads@MSNs (orange) and Pt(IV)-cov@MSNs (blue), and control MCM41-calc@MSNs (grey) at maximum concentration of 25 µM; viability measured by MTS assay after 7 days. Western Blot analysis of apoptotic/DNA damage markers for cell lines OSCC37 (**f**) and OSCC55 (**h**) after 4 days of IC75 and IC90 (3 and 4 µM respectively) concentration treatments with drug-loaded MSNs. Control MSNs (MCM41-calc@MSNs) were added at the highest (IC90) concentration.

To test the potential improvement in treatment response with MSNs delivery we treated OSCC37 and OSCC55 2D cultures with increasing concentrations of MSN-encapsulated drugs. In line with the previous data, encapsulation of Pt(IV)-biSi-2 within MSNs again showcased increased cytotoxicity against its own free drug performance and even against cisPt-ads@MSNs, with lower IC50 values for Pt(IV)-cov@MSNs (**Fig. 5E, G**). Western blot analysis of OSCC37 and OSCC55 2D treated cultures (**Fig. 5F, H**) demonstrated that OSCC37 cells endured greater DNA damage (p-γH2AX) and P53 activation after Pt(IV)-cov@MSNs treatment, resulting in visible caspase-3 cleavage at 4 µM (**Fig. 5F**). OSCC55 cells however, displayed higher P53 activation and DNA damage at 3 µM concentrations, and showing higher caspase-3 cleavage upon treatment with 4 µM Pt(IV)-cov@MSNs.

Subsequently, these resistant primary cells were grown into spheroid (**Fig. 6A,C**) and organoid models (**Fig. 6B,D**) and treated with both free drug and encapsulated cisplatin and Pt(IV)-biSi-2 before downstream processing. Both models show a delay in growth (**Fig. 6A-D**) and a slight decrease in viability (**Fig. 6A’-D’**) in response to cisplatin when compared to control conditions, which was more evident in OSCC55. Free Pt(IV)-biSi-2, had no significant effect on spheroid nor organoid size, in neither resistant-OSCC lines. Opposing to what was seen in sensitive cells, cisPt-ads@MSNs treatment improved free-cisplatin cytotoxic effect in both 3D-models, and Pt(IV)-cov@MSNs treatment demonstrated an even greater antitumoral effect, decreasing spheroid and organoid size significantly below the starting point of the experiment.

**Fig 6.**
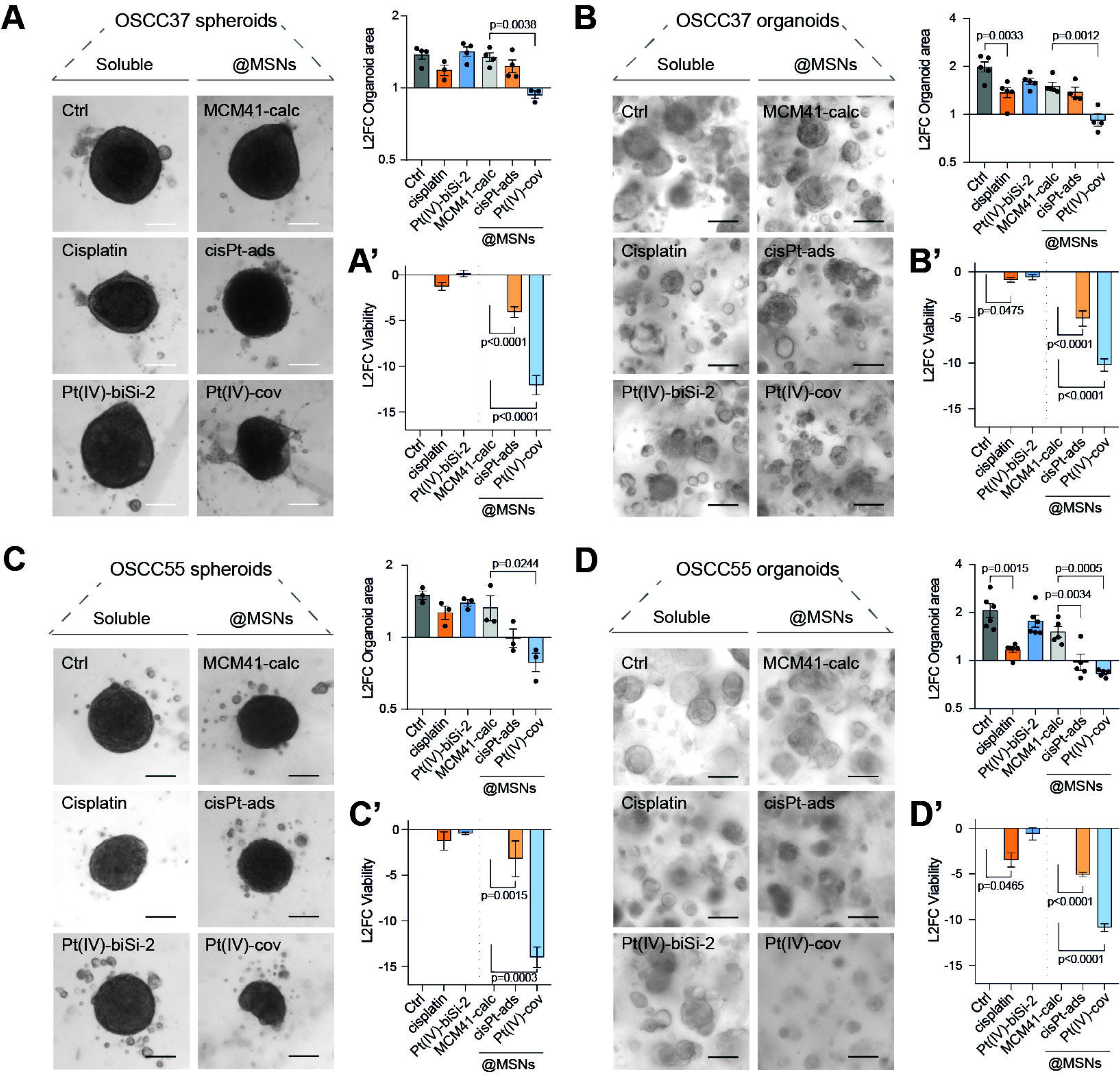
Treatment with Pt(IV)-cov@MSNs impairs cell viability in cisplatin-resistant OSCC 3D models. **a, b)** 3D *in vitro* models of cisplatin-resistant samples OSCC37 and OSCC55 were grown for 3 days before treatment with 10 µM of every drug formulation and maintained for 7 days. Left panel, representative brightfield microscopy images illustrate spheroids (**a**, **c**) and organoids (**b**, **d**) at treatment endpoint. Right panels, Graph deciphering the relative growth of the models after the treatments expressed as Log2 of fold change in area at day 7 vs. day 3. (**a’**, **b’**, **c’**, **d’**) Graphs showing changes in viability measured by Cell Glow assay and expressed as Log2 fold change of relative luminescence units (RLU) at endpoint (7 days). mean ± SEM, n = at least 3. Scale bar = 100 µm.

Since these primary cell lines form highly keratinized cores within spheroids and organoids (**Fig. 2B,C**), which difficult its processing to single cell suspension to measure apoptosis, instead, we analyzed cell viability by measuring cellular ATP levels (**Fig. 6A’-D’**). Remarkably, Pt(IV)-cov@MSNs treatment impaired spheroid and organoid growth to nearly undetectable levels of ATP. Meanwhile, cisPt-ads@MSNs elicit comparable cytotoxic responses in the OSCC55 line, which remained modest in the OSCC37 cell line model. This analysis confirmed the increased antitumoral performance of drug-loaded MSNs in comparison to the free drugs and reinforces the effect of Pt(IV)-cov@MSNs treatment on cisplatin-resistant cells.

## Discussion

This study aims to test the efficacy of the new Pt(IV)-biSi-2 prodrug and novel delivery strategies through MSNs using primary-patient derived cell cultures as a screening platform to improve the response to platinum of oral squamous carcinoma patients. 3D *in vitro* model implementation, particularly organoids[33,39] or spheroids grown within biocompatible matrices represent a relevant and rapidly expanding field that requires highly optimized protocols for downstream analysis and technical manipulation. Here, we used several methods and protocols for assessing cytotoxicity, such as flow cytometry or immunostaining, depending on the technical challenges inherent to inter-patient heterogeneity within samples. Previous work determined Pt(IV)-biSi-2 outperformed cisplatin in human intestinal tumor cell lines[35], although dose-response experiments against established and primary OSCC cell lines showed opposite results. This may be due to several factors, including uptake, GSH/AsA-dependent reduction rate and cell-intrinsic cisplatin resistance and efflux mechanisms, which can make Pt(IV) compound efficacy and cytotoxicity highly heterogeneous. Furthermore, Pt(IV)-biSi-2 was determined to be highly lipophilic, which should improve passive diffusion. This might prove true when in contact with the cell membrane but might interfere with proper dispersion and distribution in aqueous cell culture medium. We were particularly interested in probing novel delivery strategies and therefore improving antitumoral efficacy against cisplatin-resistant samples. By encapsulating Pt(IV)-biSi-2 in MSNs, unspecific binding and premature degradation are prevented, providing a focalized and targeted delivery, determined by pH acidification. Pt(IV)-cov@MSNs consistently elicited the highest cytotoxic effect in both cisplatin-sensitive and resistant primary cell lines, in both 2D and 3D models. This was relatively unexpected, since Pt(IV)-biSi-2 is covalently attached to the nanoparticles and therefore modestly released into cell at the endpoint of our experiments. Although we have shown that both free Pt(IV)-biSi-2 and Pt(IV)-cov-@MSNs induce caspase-3 - dependent apoptosis, the exact mechanism by which these nanoparticles enhance cytotoxicity to such an extent and/or whether alternative or additional stress response pathways are being activated remains to be further investigated. Despite having previously demonstrated that Pt(IV)-biSi-2 treatment produced p53-independent cytotoxicity in human intestinal tumor cell line HCT-116, cell response to DNA damage caused by Pt(IV)-biSi-2 might vary between cell type or be cancer-specific[40]. Previous work has also shown that reduction of Pt(IV)-biSi-2 resulted in the formation of the aquo Pt(II) counterparts which are the bioactive species that react with the DNA[35]. Other potential alternative source of stress might come from axial ligand composition, since these are the first moieties to be released during the reduction step[41–43]. We hypothesize that MSN encapsulation of cisplatin and Pt(IV)-biSi-2 could effectively protect against premature degradation and effectively deliver a targeted and focal release of the loaded compound. We observed that after treatment, MSNs tended to aggregate around organoids and were also visible in clusters within them (**Fig. S1A**). However, the significant increase in cytotoxicity displayed by loaded MSNs is unlikely to be due to membrane disruption or cellular stress caused by nanoparticle aggregation, since equal concentrations of vehicle (empty) nanoparticles and same levels of aggregation did not affect viability nor organoid growth.

Overall, we observe sensibilization of previously cisplatin-resistant primary OSCC cell lines in 2D and 3D *in vitro* models using the Pt(IV)-cov@MSNs. These promising results showcase the potential translational implementation of nanocarrier-based cytotoxic drug delivery to overcome important limitations associated with platinum-based systemic chemotherapy. Among these, inherent or acquired resistance to cisplatin is possibly one of the most concerning, as it inevitably compromises treatment options and patient prognosis[12,44]. Given the vast opportunities for structure modification and functionalization, nanocarrier-based drug delivery is an active field of research and development that has already shown important advances in *in vitro* and *in vivo* settings in the OSCC context[45–47]. Collectively, our results present biologically relevant drug testing models based on patient-derived tumor 3D models that recapitulate cisplatin response heterogeneity within the OSCC context. This allowed the characterization of the cytotoxic response of the Pt(IV)-biSi-2 and their respective MSN-loaded formulations. This study reiterates and highlights the need for novel approaches to target cisplatin-refractory samples, where underperforming cytotoxic drugs with better tolerance profiles can be repurposed via nanocarrier delivery for improved performance.

## DECLARATIONS

### Authors’ Contributions

Ana Belén Griso-Acevedo: validation, investigation, writing-original draft. Francisco Navas: validation, investigation, writing-original draft. Natalia Calvo: investigation. Victoria Morales: supervision, writing—review and editing. Beatriz Castelo: funding acquisition, writing—review and editing. Javier González Martín-Moro: investigation. Maria Jose Moran-Soto: investigation. Raúl Sanz: supervision, funding acquisition, writing—review and editing. José Luis Cebrián-Carretero: funding acquisition, writing—review and editing. Rafael A. García-Muñoz: conceptualization, supervision, funding acquisition, writing—review and editing. Ana Sastre-Perona: conceptualization, supervision, funding acquisition, writing-original draft, writing—review and editing.

### Consent for publication

Not applicable.

---

**Fig S1.** (**a**) Hematoxylin-Eosin staining of OSCC43 (upper row) and OSCC37 (bottom row) organoids at treatment endpoint. Arrowheads show MSN aggregates near and within organoids (**b**) Flow cytometry cell cycle analysis for Hoescht-stained single cell suspension of OSCC43 3D models: spheroids (left panel) and organoids (right panel). Scale bar = 100 µm.

## Bibliography

1 Bray F, Laversanne M, Sung H, et al. Global cancer statistics 2022: GLOBOCAN estimates of incidence and mortality worldwide for 36 cancers in 185 countries. CA: A Cancer J Clin Published Online First: 2024. [PMID: 38572751 DOI: 10.3322/caac.21834]

2 Tan Y, Wang Z, Xu M, et al. Oral squamous cell carcinomas: state of the field and emerging directions. Int J Oral Sci 2023;15:44. [PMID: 37736748 DOI: 10.1038/s41368-023-00249-w]

3 Kowalski LP, Carvalho AL, Priante AVM, Magrin J. Predictive factors for distant metastasis from oral and oropharyngeal squamous cell carcinoma. Oral Oncol 2005;41:534–41. [PMID: 15878760 DOI: 10.1016/j.oraloncology.2005.01.012]

4 Spoerl S, Gerken M, Mamilos A, et al. Lymph node ratio as a predictor for outcome in oral squamous cell carcinoma: a multicenter population-based cohort study. Clin Oral Investig 2021;25:1705–13. [PMID: 32754787 DOI: 10.1007/s00784-020-03471-6]

5 Tomioka H, Yamagata Y, Oikawa Y, et al. Risk factors for distant metastasis in locoregionally controlled oral squamous cell carcinoma: a retrospective study. Sci Rep 2021;11:5213. [PMID: 33664318 DOI: 10.1038/s41598-021-84704-w]

6 Mohamad I, Glaun MDE, Prabhash K, et al. Current Treatment Strategies and Risk Stratification for Oral Carcinoma. Am Soc Clin Oncol Educ Book 2023;43:e389810. [PMID: 37200591 DOI: 10.1200/edbk_389810]

7 Cheng Y, Li S, Gao L, Zhi K, Ren W. The Molecular Basis and Therapeutic Aspects of Cisplatin Resistance in Oral Squamous Cell Carcinoma. Front Oncol 2021;11:761379. [PMID: 34746001 DOI: 10.3389/fonc.2021.761379]

8 Elmorsy EA, Saber S, Hamad RS, et al. Advances in understanding cisplatin-induced toxicity: Molecular mechanisms and protective strategies. Eur J Pharm Sci 2024;203:106939. [PMID: 39423903 DOI: 10.1016/j.ejps.2024.106939]

9 Eljack ND, Ma H-YM, Drucker J, et al. Mechanisms of cell uptake and toxicity of the anticancer drug cisplatin. Metallomics 2014;6:2126–33. [PMID: 25306996 DOI: 10.1039/c4mt00238e]

10 Wang D, Lippard SJ. Cellular processing of platinum anticancer drugs. Nat Rev Drug Discov 2005;4:307– 20. [PMID: 15789122 DOI: 10.1038/nrd1691]

11 Siddik ZH. Cisplatin: mode of cytotoxic action and molecular basis of resistance. Oncogene 2003;22:7265–79. [PMID: 14576837 DOI: 10.1038/sj.onc.1206933]

12 Griso AB, Acero-Riaguas L, Castelo B, Cebrián-Carretero JL, Sastre-Perona A. Mechanisms of Cisplatin Resistance in HPV Negative Head and Neck Squamous Cell Carcinomas. Cells 2022;11:561. [PMID: 35159370 DOI: 10.3390/cells11030561]

13 Pendleton KP, Grandis JR. Cisplatin-Based Chemotherapy Options for Recurrent and/or Metastatic Squamous Cell Cancer of the Head and Neck. Clin Medicine Insights Ther 2013;5:CMT.S10409. [PMID: 24273416 DOI: 10.4137/cmt.s10409]

14 Specht L, Larsen SK, Hansen HS. Phase II study of docetaxel and cisplatin in patients with recurrent or disseminated squamous-cell carcinoma of the head and neck. Ann Oncol 2000;11:845–9. [PMID: 10997812 DOI: 10.1023/a:1008355315205]

15 Štarha P, Křikavová R. Platinum(IV) and platinum(II) anticancer complexes with biologically active releasable ligands. Coord Chem Rev 2024;501:215578. [DOI: 10.1016/j.ccr.2023.215578]

16 Xu L, Kong X, Li X, et al. Current Status of Novel Multifunctional Targeted Pt(IV) Compounds and Their Reductive Release Properties. Molecules 2024;29:746. [PMID: 38398498 DOI: 10.3390/molecules29040746]

17 Jia C, Deacon GB, Zhang Y, Gao C. Platinum(IV) antitumor complexes and their nano-drug delivery. Coord Chem Rev 2021;429:213640. [DOI: 10.1016/j.ccr.2020.213640]

18 Shahlaei M, Asl SM, Derakhshani A, et al. Platinum-based drugs in cancer treatment: Expanding horizons and overcoming resistance. J Mol Struct 2024;1301:137366. [DOI: 10.1016/j.molstruc.2023.137366]

19 Zhang C, Xu C, Gao X, Yao Q. Platinum-based drugs for cancer therapy and anti-tumor strategies. Theranostics 2022;12:2115–32. [PMID: 35265202 DOI: 10.7150/thno.69424]

20 Johnstone TC, Suntharalingam K, Lippard SJ. The Next Generation of Platinum Drugs: Targeted Pt(II) Agents, Nanoparticle Delivery, and Pt(IV) Prodrugs. Chem Rev 2016;116:3436–86. [PMID: 26865551 DOI: 10.1021/acs.chemrev.5b00597]

21 Zhang S, Wang X, Guo Z. Rational design of anticancer platinum(IV) prodrugs. Adv Inorg Chem 2019;149–82. [DOI: 10.1016/bs.adioch.2019.10.009]

22 Hall MD, Hambley TW. Platinum(IV) antitumour compounds: their bioinorganic chemistry. Coord Chem Rev 2002;232:49–67. [DOI: 10.1016/s0010-8545(02)00026-7]

23 Yempala T, Babu T, Karmakar S, et al. Expanding the Arsenal of PtIV Anticancer Agents: Multi-action PtIV Anticancer Agents with Bioactive Ligands Possessing a Hydroxy Functional Group. Angew Chem Int Ed 2019;58:18218–23. [PMID: 31599054 DOI: 10.1002/anie.201910014]

24 Guo H, Zhao X, Duan Y, Shi J. Hollow mesoporous silica nanoparticles for drug formulation and delivery: Opportunities for cancer therapy. Colloids Surf B: Biointerfaces 2025;249:114534. [PMID: 39874869 DOI: 10.1016/j.colsurfb.2025.114534]

25 Truong-Thi N-H, Nguyen NH, Nguyen DTD, Tang TN, Nguyen TH, Nguyen DH. pH-responsive delivery of Platinum-based drugs through the surface modification of heparin on mesoporous silica nanoparticles. Eur Polym J 2023;185:111818. [DOI: 10.1016/j.eurpolymj.2023.111818]

26 Xu B, Li S, Shi R, Liu H. Multifunctional mesoporous silica nanoparticles for biomedical applications. Signal Transduct Target Ther 2023;8:435. [PMID: 37996406 DOI: 10.1038/s41392-023-01654-7]

27 Fatima R, Katiyar P, Kushwaha K. Recent advances in mesoporous silica nanoparticle: synthesis, drug loading, release mechanisms, and diverse applications. Front Nanotechnol 2025;7:1564188. [DOI: 10.3389/fnano.2025.1564188]

28 Morales V, Gutiérrez-Salmerón M, Balabasquer M, et al. New Drug-Structure-Directing Agent Concept: Inherent Pharmacological Activity Combined with Templating Solid and Hollow-Shell Mesostructured Silica Nanoparticles. Adv Funct Mater 2016;26:7291–303. [DOI: 10.1002/adfm.201505073]

29 Pérez-Garnes M, Gutiérrez-Salmerón M, Morales V, et al. Engineering hollow mesoporous silica nanoparticles to increase cytotoxicity. Mater Sci Eng: C 2020;112:110935. [PMID: 32409082 DOI: 10.1016/j.msec.2020.110935]

30 Morales V, Idso MN, Balabasquer M, Chmelka B, García-Muñoz RA. Correlating Surface-Functionalization of Mesoporous Silica with Adsorption and Release of Pharmaceutical Guest Species. J Phys Chem C 2016;120:16887–98. [DOI: 10.1021/acs.jpcc.6b06238]

31 García-Muñoz RA, Morales V, Linares M, González PE, Sanz R, Serrano DP. Influence of the structural and textural properties of ordered mesoporous materials and hierarchical zeolitic supports on the controlled release of methylprednisolone hemisuccinate. J Mater Chem B 2014;2:7996–8004. [PMID: 32262090 DOI: 10.1039/c4tb00089g]

32 He H, Xiao H, Kuang H, et al. Synthesis of mesoporous silica nanoparticle–oxaliplatin conjugates for improved anticancer drug delivery. Colloids Surf B: Biointerfaces 2014;117:75–81. [PMID: 24632033 DOI: 10.1016/j.colsurfb.2014.02.014]

33 Chaves P, Garrido M, Oliver J, Pérez-Ruiz E, Barragan I, Rueda-Domínguez A. Preclinical models in head and neck squamous cell carcinoma. Br J Cancer 2023;128:1819–27. [PMID: 36765175 DOI: 10.1038/s41416-023-02186-1]

34 Driehuis E, Kolders S, Spelier S, et al. Oral Mucosal Organoids as a Potential Platform for Personalized Cancer Therapy. Cancer Discov 2019;9:852–71. [PMID: 31053628 DOI: 10.1158/2159-8290.cd-18-1522]

35 Navas F, Chocarro-Calvo A, Iglesias-Hernández P, et al. Promising Anticancer Prodrugs Based on Pt(IV) Complexes with Bis-organosilane Ligands in Axial Positions. J Med Chem 2024;67:6410–24. [PMID: 38592014 DOI: 10.1021/acs.jmedchem.3c02393]

36 Schober M, Fuchs E. Tumor-initiating stem cells of squamous cell carcinomas and their control by TGF-β and integrin/focal adhesion kinase (FAK) signaling. Proc National Acad Sci 2011;108:10544–9. [PMID: 21670270 DOI: 10.1073/pnas.1107807108]

37 Berg S, Kutra D, Kroeger T, et al. ilastik: interactive machine learning for (bio)image analysis. Nat Methods 2019;16:1226–32. [PMID: 31570887 DOI: 10.1038/s41592-019-0582-9]

38 Driehuis E, Kretzschmar K, Clevers H. Establishment of patient-derived cancer organoids for drug-screening applications. Nat Protoc 2020;15:3380–409. [PMID: 32929210 DOI: 10.1038/s41596-020-0379-4]

39 Warren A, Chen Y, Jones A, et al. Global computational alignment of tumor and cell line transcriptional profiles. Nat Commun 2021;12:22. [PMID: 33397959 DOI: 10.1038/s41467-020-20294-x]

40 Zhao C-L, Qiao X, Liu X-M, et al. Rapid DNA interstrand cross-linking of Pt(IV) compound. Eur J Pharmacol 2022;925:174985. [PMID: 35489419 DOI: 10.1016/j.ejphar.2022.174985]

41 Tolan D, Almotairy ARZ, Howe O, Devereux M, Montagner D, Erxleben A. Cytotoxicity and ROS production of novel Pt(IV) oxaliplatin derivatives with indole propionic acid. Inorg Chim Acta 2019;492:262–7. [DOI: 10.1016/j.ica.2019.04.038]

42 Tolan D, Gandin V, Morrison L, et al. Oxidative Stress Induced by Pt(IV) Pro-drugs Based on the Cisplatin Scaffold and Indole Carboxylic Acids in Axial Position. Sci Rep 2016;6:29367. [PMID: 27404565 DOI: 10.1038/srep29367]

43 Tabrizi L, Jones AM, Romero-Canelon I, Erxleben A. Multiaction Pt(IV) Complexes: Cytotoxicity in Ovarian Cancer Cell Lines and Mechanistic Studies. Inorg Chem 2024;63:14958–68. [PMID: 39083592 DOI: 10.1021/acs.inorgchem.4c01586]

44 Chen S-H, Chang J-Y. New Insights into Mechanisms of Cisplatin Resistance: From Tumor Cell to Microenvironment. Int J Mol Sci 2019;20:4136. [PMID: 31450627 DOI: 10.3390/ijms20174136]

45 Elsaady SA, Aboushelib MN, Al-Wakeel E, Badawi MF. A novel intra-tumoral drug delivery carrier for treatment of oral squamous cell carcinoma. Sci Rep 2023;13:11984. [PMID: 37491569 DOI: 10.1038/s41598-023-38230-6]

46 Li R, Wan C, Li Y, et al. Nanocarrier-based drug delivery system with dual targeting and NIR/pH response for synergistic treatment of oral squamous cell carcinoma. Colloids Surf B: Biointerfaces 2024;244:114179. [PMID: 39217727 DOI: 10.1016/j.colsurfb.2024.114179]

47 Cui S, Liu H, Cui G. Nanoparticles as drug delivery systems in the treatment of oral squamous cell carcinoma: current status and recent progression. Front Pharmacol 2023;14:1176422. [PMID: 37292147 DOI: 10.3389/fphar.2023.1176422]

