## Supplementary figures and images for "Overcoming Cisplatin Resistance in 3D Oral Squamous Cell Carcinoma Models via Nanoparticle-Mediated Pt(IV) Drug Delivery"

### Figure S1

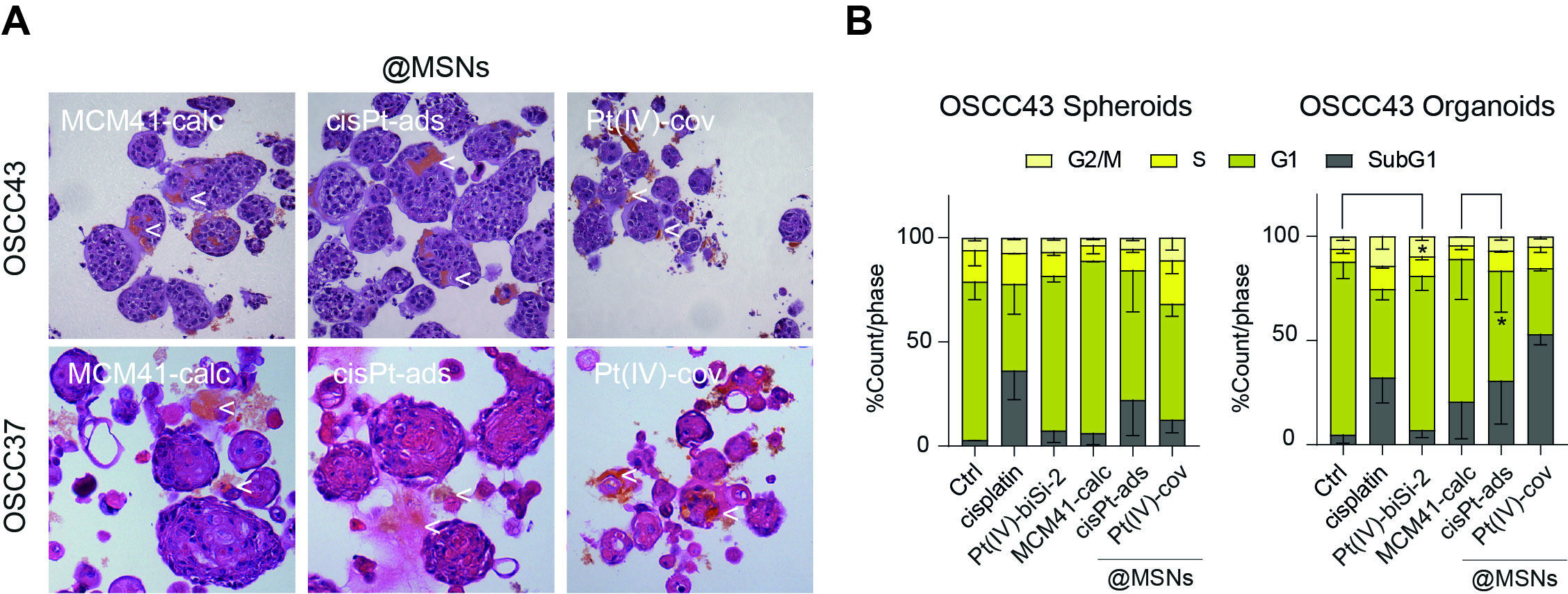
